# Movi: a fast and cache-efficient full-text pangenome index

**DOI:** 10.1101/2023.11.04.565615

**Authors:** Mohsen Zakeri, Nathaniel K. Brown, Omar Y. Ahmed, Travis Gagie, Ben Langmead

## Abstract

Efficient pangenome indexes are promising tools for many applications, including rapid classification of nanopore sequencing reads. Recently, a compressed-index data structure called the “move structure” was proposed as an alternative to other BWT-based indexes like the FM index and r-index. The move structure uniquely achieves both O(r) space and O(1)-time queries, where r is the number of runs in the pangenome BWT. We implemented Movi, an efficient tool for building and querying move-structure pangenome indexes. While the size of the Movi’s index is larger than the r-index, it scales at a smaller rate for pangenome references, as its size is exactly proportional to r, the number of runs in the BWT of the reference. Movi can compute sophisticated matching queries needed for classification – such as pseudo-matching lengths and backward search – at least ten times faster than the fastest available methods, and in some cases more than 30-fold faster. Movi achieves this speed by leveraging the move structure’s strong locality of reference, incurring close to the minimum possible number of cache misses for queries against large pangenomes. We achieve still further speed improvements by using memory prefetching to attain a degree of latency hiding that would be difficult with other index structures like the r-index. Movi’s fast constant-time query loop makes it well suited to real-time applications like adaptive sampling for nanopore sequencing, where decisions must be made in a small and predictable time interval.

## 1 Introduction

Pangenome indexes are promising tools for aligning and classifying sequencing reads with respect to large sets of similar reference sequences. While many existing tools are k-mer based [1, 2], others use flexible indexes enabling arbitrary-length pattern matching queries, like the `FM`-index [3, 4] and r-index [5, 6]. The `FM`-index [7] and r-index [8] are full-text indexes that facilitate matching via “backward search.” The r-index can also find maximal exact matches (MEMs) and matching statistics using the MONI algorithm [9]. Unlike the `FM`-index, the r-index is run-length compressed, allowing the index to grow proportionally to the amount of *distinct* sequence in a pangenome reference, rather than its total length.

In practice, the r-index comprises a collection of data structures such as bitvectors and wavelet tries. A single query – such as a backward-search step – involves memory accesses to many disparate memory addresses within these structures. The number and unpredictability of these accesses leads to cache misses, i.e. pauses during which the processor is stalled waiting for portions of the data structures to be moved from main memory to nearby cache memories. Even when the time required for an index query is theoretically constant, the latency incurred by cache misses can be large, making queries slow in practice. Variability in the number of cache misses incurred per query leads to fluctuating latency across queries. Overall, the effect is to make queries slow with high variability.

The Move structure was introduced by Nishimoto and Tabei in 2021 [10]. Like the `FM`-index and r-index, it is a full-text index based on the Burrows Wheeler Transform (BWT). It achieves both O(r) space usage and O(1) (constant) time for LF mapping queries. This combination has not been achieved by other indexes; e.g. the r-index can achieve one or the other but not both. Another key advantage of the move structure is that it consists entirely of a single table. Move structure queries need only perform a limited number of accesses to this table, incurring few – usually just one or two – cache misses per query. That is, move structure queries have excellent locality of reference. This leads to faster queries with more predictable latency compared to alternatives like the r-index. While past studies have shown some of the move structure’s computational trade-offs relative to r-index [11], no studies have investigated these advantages related to speed and locality of reference.

Here we introduce Movi, a pangenome full-text index based on the move structure. Movi is much faster than alternative pangenome indexes like the r-index. We measure Movi’s cache characteristics and show that queries achieve a small, nearly-minimal number of cache misses. Further, we show that the latency of the remaining cache misses can be “hidden” to a large degree by rearranging the computation and using memory prefetch instructions. We demonstrate that Movi can implement the same algorithms as alternative pangenome tools like r-index (backward search) and SPUMONI (pseudo-matching lengths and matching statistics), while running drastically faster, e.g. 30 times faster than SPUMONI. Finally, we show that despite having a larger size compared to other pangenome indexes, Movi’s index grows more slowly than other pangenome indexes as genomes are added.

In short, Movi is the fastest available tool for full-text pangenome indexing and querying, and our open source implementation enables its application in various classification and alignment scenarios, including in speed-critical scenarios like adaptive sampling for nanopore sequencing.

## 2 Methods

### 2.1 Burrows Wheeler Transform, FM-index and r-index

The Burrows Wheeler Transform (BWT) is a reversible permutation that reorders the characters of a string T according to the lexicographical order of their right contexts in T. Beginning with T of length n, we append a terminal symbol $ that does not appear elsewhere in T and is lexicographically smaller than T’s other characters. T[i] denotes the character at 1-based offset i and T[i..n] denotes a suffix starting at i. BWT(T) permutes T’s characters so that T[i] comes before T[j] in BWT order if and only if T[i+1 .. n] *<* T[j+1 .. n]. Repetitive portions of T yield long “runs” in BWT(T) where a run is a maximal-length substring consisting of a character repeated. Figure 1a illustrates BWT runs of length up to 8 in the last column of the matrix.

**Figure 1:**
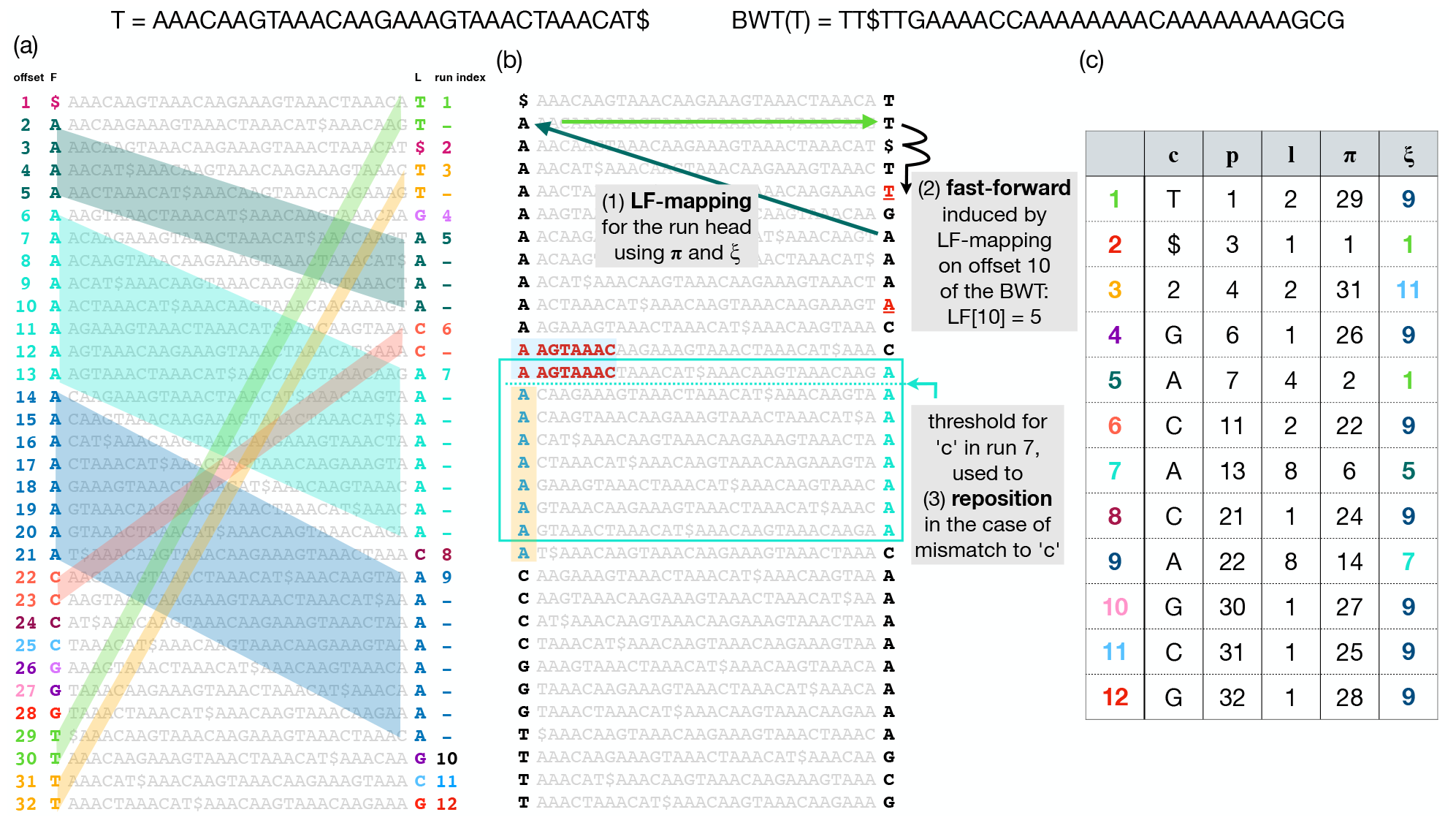
Top: T and BWT(T). (a) BWM(T), consisting of T’s sorted rotations. The leftmost column is called F and the rightmost column is BWT(T), also called L. Distinct BWT runs are given distinct colors. The `LF`-mapping maps these runs to same-letter stretches in F. This is illustrated using matching colors and, in the case of multi-character runs, by parallelograms connecting BWT characters to their counterparts in F. (b) Arrows at the top illustrate how a move-structure query for LF[10] results in one LF step (green arrows) followed by two fast forward steps (black arrow). Below, the blue arrows illustrate how a threshold facilitates “repositioning.” A a mismatch between the BWT character (“A”) and a “C” from the query causes a jump to the nearest offset above or below ending in “C.” The one above was chosen in this case because it has a longer longest common prefix (LCP) with the rotation at the original offset. The threshold (blue dotted line) denotes the point above which rows have a longer LCP with the next C-terminated row above, but rows below have a longer LCP with the next C-terminated row below (with ties broken arbitrarily). (c) Each BWTrun is a row in the move structure table; c is the run character, *ℓ* is the length, p is the offset with respect to the BWT, π is LF[p], and ξ is the index of the run containing offset π.

Figure 1a and Figure 1b show two copies of a Burrows-Wheeler Matrix or BWM. Rows of the BWM consist of all distinct rotations of the string T ordered lexicographically. BWT(T) is the last column of BWM(T). BWM’s first and last columns are related by the Last-to-First mapping (“`LF`-mapping”) [7], which states that the i^th^ occurrence of a character c in the last column of the BWM corresponds to the same text occurrence as the i^th^ occurrence of c in the first column. Some `LF`-mapping relationships are illustrated with parallelograms in Figure 1a. The `LF`-mapping also gives a way to navigate through the text T. Note that if the BWT permutation maps T[j] to BWT[i], then LF[i] gives the BWT index of T[j – 1] (or T[n] if j = 1). So the `LF`-mapping allows for right-to-left movements with respect to T, a fact used in pattern-matching queries.

The FM-index is a data structure based on BWT(T) enabling fast and efficient computation of the `LF`-mapping and related queries. It consists of BWT(T) as well as succinct data structures for storing and querying character ranks within BWT(T). In typical implementations, it grows linearly with the text: O(n).

When T is repetitive, the number of BWT runs (`r`) is much smaller than the text length (n). The r-index [8] exploits this by representing the BWT in a run-length-compressed fashion. This version is called the RLBWT. The i^th^ run, denoted RLBWT[i], consists of the character repeated in the run (RLBWT[i].c), and the run’s length (RLBWT[i].n). Additional data structures enable efficient computation of the `LF`-mapping without having to decompress the RLBWT. The data structures making up the r-index fit in O(`r`) space total.

### 2.2 The Move Data Structure

#### LF Mapping

When T is a repetitive pangenome, the `LF`-mapping tends to map consecutive stretches of BWT characters to consecutive stretches in F (Figure 1a). The move structure exploits this to simplify computation of the `LF`-mapping. The move structure consists of a table (M) with rows corresponding to BWT runs (Figure 1c). To aid `LF`-mapping, the column named π stores the `LF`-mapping of the run head, i.e. M[i].π = LF[M[i].p]. To compute the `LF`-mapping at any offset in run index i, we begin by following M[i].π. This will either jump to the correct run, or to a run preceding the correct one.

Given M, a BWToffset j, and a run index i, we compute LF[j] by adding j’s offset into the current run (j – M[i].p) to the run head’s `LF`-mapping:

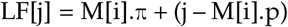

Note that this involves only arithmetic on i, j, M[i].π and M[i].p, and does not involve bitvectors or wavelet-tree queries. Only the accesses to M[i] might require accessing main memory. An illustration of how memory accesses induced by move structure queries differ than those by r-index is shown in Figure S4.

Note that an input to this computation is i, the current run index. To chain multiple `LF`-mapping queries together, as is needed for matching queries, we must update not only the BWT offset j but also the BWT run index i. As a step toward this goal, M[i].ξ stores the index of the run containing LF[M[i].π]. However, the run containing LF[M[i].π] may not also contain LF[j]. I.e. it is possible that LF[j] – M[M[i].ξ].p *>* M[M[i].ξ].*C*. After jumping to M[M[i].ξ], we may additionally need to advance through the runs until finding the smallest run index i^*t*^ *>* i such that M[i^*t*^].p *≤* LF[j] *<* M[i^*t*^ + 1].p. We call this the “fast-forward” or “ff” procedure, illustrated in Figure 1b (top) and detailed by Algorithm S1 in supplementary materials. Using that algorithm, we update both i and j in each `LF`-mapping step:

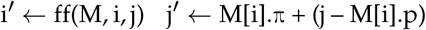

#### Constant-time LF

Nishimoto and Tabei gave a procedure for splitting some BWT runs into shorter sub-runs to achieve a constant upper bound on the fast-forwards required for any `LF`-mapping [10]. The procedure works with a parameter d such that, after splitting runs, the number of fast-forwards per `LF`-mapping query is less than 2d while adding at most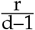 additional runs to the table. The overall number of runs is still O(r) after splitting. In practice, the procedure splits only a fraction of the original runs [12].

With the exception of the jump induced by following LF[M[i].π], all the memory accesses described here are sequential. The LF[M[i].π] step is unpredictable, possibly needing to access a distant and not-recently-accessed location in memory, likely incurring a cache miss. That said, for a chain of several `LF`-mapping queries, only one expensive memory access is needed per query.

Since information about exact BWT offsets of matches is not required for computing pseudo matching lengths, Movi avoids storing both p and π in the table. Instead, Movi collapses those fields into a single relative offset, as previously implemented by Brown et al [11].

### 2.3 Computing pseudo matching lengths

#### Matching statistics and pseudo matching lengths

Matching statistics (MS) are a summary of sequence similarity used in sequence classification tasks and for computing other similarity features like Maximal Exact Matches (MEMs). Given a text T[1..n] and pattern P[1..m], P’s matching statistics with respect to T are defined as an array MS[1..m] where MS[i] is the length of the longest prefix of P[i..m] occurring in T.

Bannai et al. [13] described a 2-pass procedure for computing matching statistics using the r-index and an auxiliary thresholds structure. Rossi et al. [9] gave an efficient procedure for computing the thresholds. Later, Ahmed et al. [5] introduced a modified 1-pass version of the procedure that computes a vector of Pseudo Matching Lengths (PMLs), which roughly approximate the lengths in MS. While PMLs contain less information than MSs – e.g. they cannot be used to exactly compute MEMs – finding PMLs is much faster, can be performed in single pass over the query, and requires neither a suffix-array sample nor a random-access structure for T. In practice, PMLs are similar to MSs in their ability to classify sequences [5].

The MONI algorithm starts at an arbitrary offset in the BWT, then considers each character of the query sequence in right-to-left order. Say we are currently at offset j in BWT and are examining character P[i].

The algorithm first tests if P[i] = BWT[j]. If they are equal, we call this “case 1.” For case 1, the algorithm performs an `LF`-mapping step and moves on to the next character:

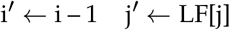

The `LF`-mapping uses the strategy we discussed above, which includes the fast-forward procedure. If P[i] ≠ BWT[j], we call this “case 2.” For case 2, we cannot simply use LF[j] as our next offset; rather we must “reposition” to a nearby offset j^r^ such that P[i] = BWT[j^r^]. We let j^r^ equal one of two choices: the greatest j^up^ such that j^up^ *<* j and BWT[j^up^] = P[i], or the smallest j^dn^ such that j^dn^ *>* j and BWT[j^dn^] = P[i]. Whether we choose j^up^ or j^dn^ is determined by querying the thresholds structure of Bannai et al. [13]. Once we have repositioned, we proceed using the same update rule as above, substituting j^r^ for j.

Instances where we can apply the simpler case 1 update rule correspond to instances where an existing match is being extended by 1 character, causing the matching statistic to increase by one. Instances where we apply case 2 might or might not correspond to an extension. The MS algorithm from MONI is capable of distinguishing these two subcases of case 2. The PML algorithm of SPUMONI is not capable of this, instead resetting the match length to 0 when it reaches an instance of case 2. Details about the PML computation procedure is shown by Algorithm S2 in supplementary materials.

#### movi’s repositioning

Movi uses two distinct strategies for finding and moving to the run containing j^r^. By default Movi, scans from run to run (either upward to j^up^ or downward to j^dn^) until reaching a run with a matching character. This involves an unpredictable number of memory accesses, though they are sequential accesses. In its Movi-constant mode (discussed further in Methods 2.5), Movi instead stores explicit pointers to the j^r^-containing runs for each characters of the DNA alphabet. It stores two such sets of pointers, one for when the threshold points upward and one for when it points downward, leading to a total of six additional pointers being stored in each move structure run.

In short, there are three types of operations performed by Movi during the PML computation; (1) jump to the run potentially containing the `LF`-mapping destination, (2) fast-forward to the run that contains the `LF`-mapping destination, and (3) reposition to the run containing a matching character in the case of a mismatch. These are illustrated in a state diagram in Figure 2 (with further detail in supplementary materials Figure S1). Operation (1) is used in each `LF`-mapping, once per base, and has the highest cost since it usually incurs a cache miss. Operations (2) and (3) are less expensive.

**Figure 2:**
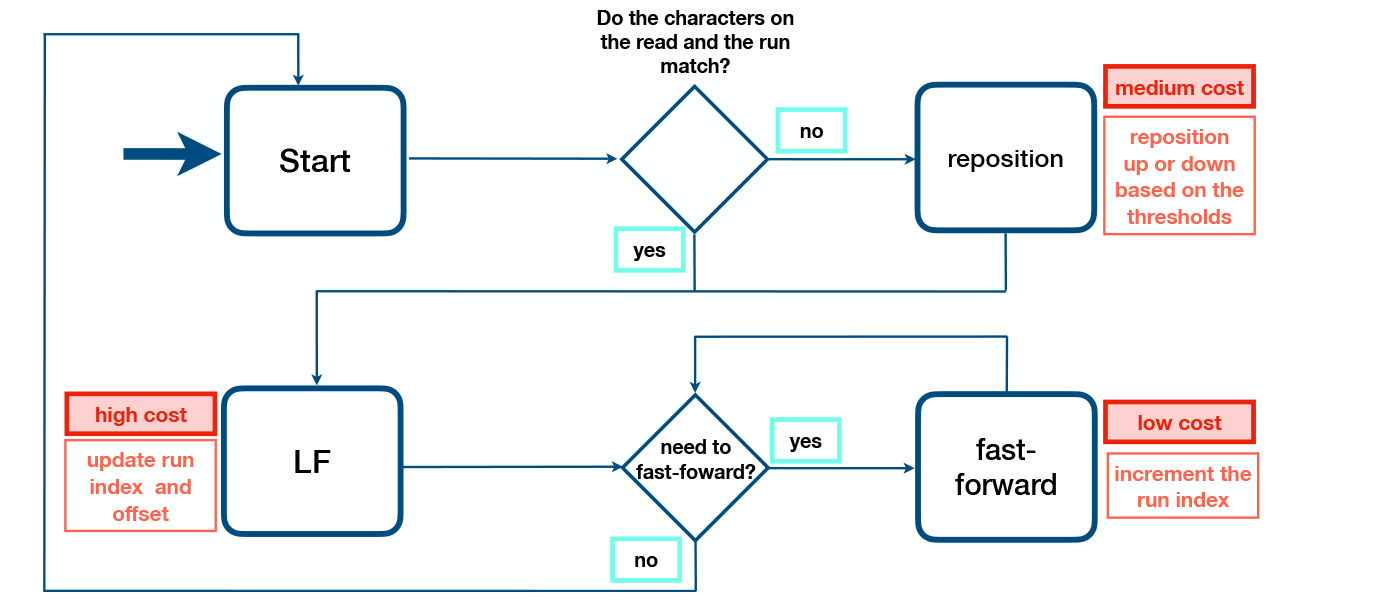
Schematic of PML computation with Movi. The typical cost associated with each memory access is shown. Higher costs are incurred by accesses that move long distances to memory addresses that have not been used recently.

### 2.4 Cache-miss latency hiding

The once-per-iteration `LF`-mapping step is Movi’s single most expensive operation. Because it moves to a new, unpredictable address, it usually incurs a cache miss and thereby stalls the processor until the requested memory (and associated cache line) is retrieved and installed in the cache.

We observed that Movi’s simple inner loop can be easily rearranged and augmented with memory “prefetch” instructions in a way that avoids the latency of stalling. To achieve this, a single thread of execution processes several reads concurrently. Say that Movi is computing PMLs for two reads named `read_a` and `read_b` concurrently in a single thread. Rather than compute all of `read_a`’s PMLs then all of `read_b`’s PMLs, Movi alternates repeatedly between the reads. It first advances the computation for `read_a` until reaching the first instruction that accesses memory in the `LF`-mapping step’s destination row. Instead of attempting the access immediately, Movi issues a memory prefetch instruction, which asynchronously requests that the needed memory be retrieved into the cache. Because this happens asynchronously, it does not immediately cause the processor to stall. Movi then switches to `read_b` and advances that computation in a similar way, ending with the prefetch of the destination row of `read_b`’s next `LF`-mapping step. Movi then switches back to `read_a`; in the meantime, the processor has at least begun (and has possibly completed) the process of installing the earlier-requested memory into the cache. We now resume the computation for `read_a`, expecting that accessing the destination row of the `LF`-mapping can now be done with little or no stalling. This process repeats until all the PMLs are computed for bot reads. We give an illustration in Figure S2 for the strategy, the pseudocode is also provided in Algorithm S3. Note that the pseudocode handles various special cases of interest, e.g. detecting when a read’s sequence has been exhausted and loading the next read.

This “latency hiding” strategy depends on a parameter: the number of reads handled concurrently. If too few reads are handled concurrently, only a fraction of the stalling time is avoided and the benefit is small. If too many reads are handled concurrently, the time between the prefetch and the actual use of the memory can become so long that competing threads and processes have caused the cache line to be erased (“evicted”) from the cache, and the stall occurs anyway. We measured the speed of Movi’s PML computation when using 2, 4, 8, 16, and 32 concurrent reads. Once reaching about 8 concurrent reads, the gain from prefetching began to plateau Figure S3. We therefore chose 16 as the default number of concurrent reads, and that setting is used in all results in the manuscript. Use of 16 concurrent reads improved throughput 2.24-fold compared to when latency hiding was disabled.

### 2.5 The Movi software

Movi supports two modes of operation. The first mode, called Movi-default, is fast but lacks the constant-time `LF`-mapping query guarantee. The second mode, called Movi-constant, uses the splitting to create a move structure that has a constant-time `LF`-mapping guarantee. Further, Movi-constant uses a constant-time version of the repositioning step, allowing its inner loop to be fully constant-time, regardless of whether it involves `LF`-mapping steps and/or repositioning. This comes at the cost of additional space, since (a) the move structure that results from the splitting procedure has more runs and is therefore somewhat larger than the unsplit move structure, and (b) the constant-time repositioning step requires that we pre-compute upward and downward jump distances and store them in the move structure table.

To build the Burrows Wheeler Transform, Movi uses the prefix-free parsing (PFP) algorithm of Boucher et al [14], which is particularly efficient for building the BWT of a highly repetitive text such as a pangenome. The algorithm also integrates Rossi et al’s [9] approach for computing thresholds for repositioning.

Movi is implemented in C++. It is GPL3-licensed open-source software available from https://github.com/mohsenzakeri/movi. It depends on both the prefix-free parsing implementation from the pfp thresholds repository ^1^ and the run splitting implementation from the r-permute library ^2^. Results in this manuscript are based on the `v1.0` tag of that repository.

## 3 Results

We measured Movi’s speed and cache characteristics relative to the related SPUMONI approach as well as to other approaches that use the FM Index (Bowtie 2), a pangenome k-mer index (Fulgor) or other approaches that achieve compression (minimap2). We measure the predictability of Movi’s innermost loop, to assess its utility for real-time data processing applications. Finally, we explores how Movi’s index scales when applied to genomes from the Human Pangenome Reference Consortium (HPRC)[15]. Experiments were run on 3 GHz Intel Xeon Gold Cascade Lake 6248R CPU with 1.5TB DDR4 memory.

### 3.1 Pseudo matching lengths for a mock community

We first measured the move structure’s efficiency for computing pseudo matching lengths (PMLs), an approximation of matching statistics previously shown to be useful for classification tasks, including adaptive sampling [5, 6]. We compared Movi’s default and constant modes to SPUMONI in terms of index size and query time. We ran the tools on the Zymo High Molecular Weight Mock Microbial Community (SRR11071395) previously used to evaluate Uncalled [16].

For further context, we also evaluated the `FM`-index based tool Bowtie2, the minimizer and hashtable-based tool minimap2, and the colored compacted de-bruijn graph-based tool Fulgor. Note that these tools differ in what they actually compute, with Bowtie2 and minimap2 generating full read alignments, and Fulgor producing pseudo-alignments. The sample consists of about 800K long reads sequenced by Oxford Nanopore Techonologies (ONT) with the average length of 15K bases.

For all tools, the index consisted of all the complete reference genomes of 7 bacteria species (Bacillus subtilis, Enterococcus faecalis, Escherichia coli, Listeria monocytogenes, Pseudomonas aeruginosa, Salmonella enterica, and Staphylococcus aureus). These were all obtained from RefSeq database [17].

Table 1 shows the size of the indexes built by all the tools as well as the time required for querying all the reads. We first compared the computational requirements of Movi-default to SPUMONI. We observed that Movi-default was 30 times faster than SPUMONI, but its index was 4.7 times larger than SPUMONI’s. Movi-constant was both slower and had a larger index compared to Movi-default; as we show later, however, the Movi-constant mode benefits from more predictable performance across inner-loop iterations.

**Table 1:** Indexes are built over all available complete genomes of 7 bacteria from RefSeq database. The size of the fasta file including the recursive complement is 67 GB. The number of long reads in the sample is 800K.^*^ The minimap2 is run with 16 threads unlike other tools which are run with a single thread. ^+^ 12x2 shows the size of two `FM`-index in the Bowtie2’s index (the forward and reverse strand). ^†^ The size of the Fulgor’s index is breakdown into two parts; the size of the k-mer set is 0.65 GB and the size of the index related to the document (color) information is 2.34 GB.

| Tool | Index type | Full-text? | Query type | Document data (color) | Size (GB) | Query time (hh:mm:ss) |
| --- | --- | --- | --- | --- | --- | --- |
| Movi-default | move structure | yes | PML | no | 8.5 | 00:18:40 |
| Movi-constant | move structure | yes | PML | no | 14 | 00:24:01 |
| SPUMONI | r-index | yes | PML | no | 1.8 | 09:20:55 |
| Bowtie2 | FM-index | yes | alignments | yes | 12x2 + 40<sup>+</sup> | - |
| Minimap2 | minimizers | no | alignments | yes | 68 | 21:31:28<sup>*</sup> |
| Fulgor | ccdbg (kmers) | no | pseudo-alignments | yes | 0.65 + 2.34<sup>†</sup> | 01:11:51 |

Fulgor had both a smaller index and a relatively fast query time compared even to Movi, taking only about 3.8 times the amount of time as Movi-default. Fulgor’s full index takes about 3 GB, about one third the size of Movi-default’s 8.5 GB index. On the other hand, the two tools output different results, with Movi outputting pseudo-matching lengths and Fulgor outputting pseudo-alignment information. Further, Fulgor is k-mer based and requires pre-selection of a set k-mer length, whereas Movi is a full-text index. Movi-default is the fastest overall and provides an advantageous trade for applications that benefit from the flexibility of a full-text index, e.g. adaptive sampling.

Bowtie2 and minimap2 are not perfectly comparable to Movi since they produce full read alignments. Further Bowtie2 is designed for use with short reads, not the long nanopore reads assessed here. For that reason, we omitted Bowtie2 from the speed comparison. Minimap2 took about 69 times longer to align the reads, while also using 16 threads (compared to 1 thread for the other tools). Its index was also 8 times larger than Movi-default’s. So although minimap2 is able to produce full and accurate alignments for the nanopore reads (Movi only computes the pseudo matching lengths), Movi provides a useful combination of speed and memory efficiency for applications, such as classification, where pseudo matching lengths provide sufficient power.

Finally, we compared the PMLs generated by Movi (both modes) against those computed by SPUMONI. Using the `diff` tool, we found that Movi and SPUMONI generated identical PMLs, as expected.

### 3.2 Speed and predictability of Movi queries

Because of its simple tabular form, we hypothesized the move structure would exhibit superior cache characteristics compared to SPUMONI. We used the “Cachegrind^3^” profiler to measure the cache misses incurred by Movi and SPUMONI when computing PMLs for the same Zymo sample used in the previous section. Specifically, we measured misses in the “last-level” cache, i.e. the final level of cache before main memory, since these are the misses that take the most time.

Figure 3a shows the number of cache misses per base. We observed that SPUMONI incurred more than 14 times as many cache misses per base compared to Movi. The reduced cache miss rate of Movi came at the cost of a larger index. We also observed that the time required for each iteration of the inner loop was both smaller and less variable for Movi compared to SPUMONI.

**Figure 3:**
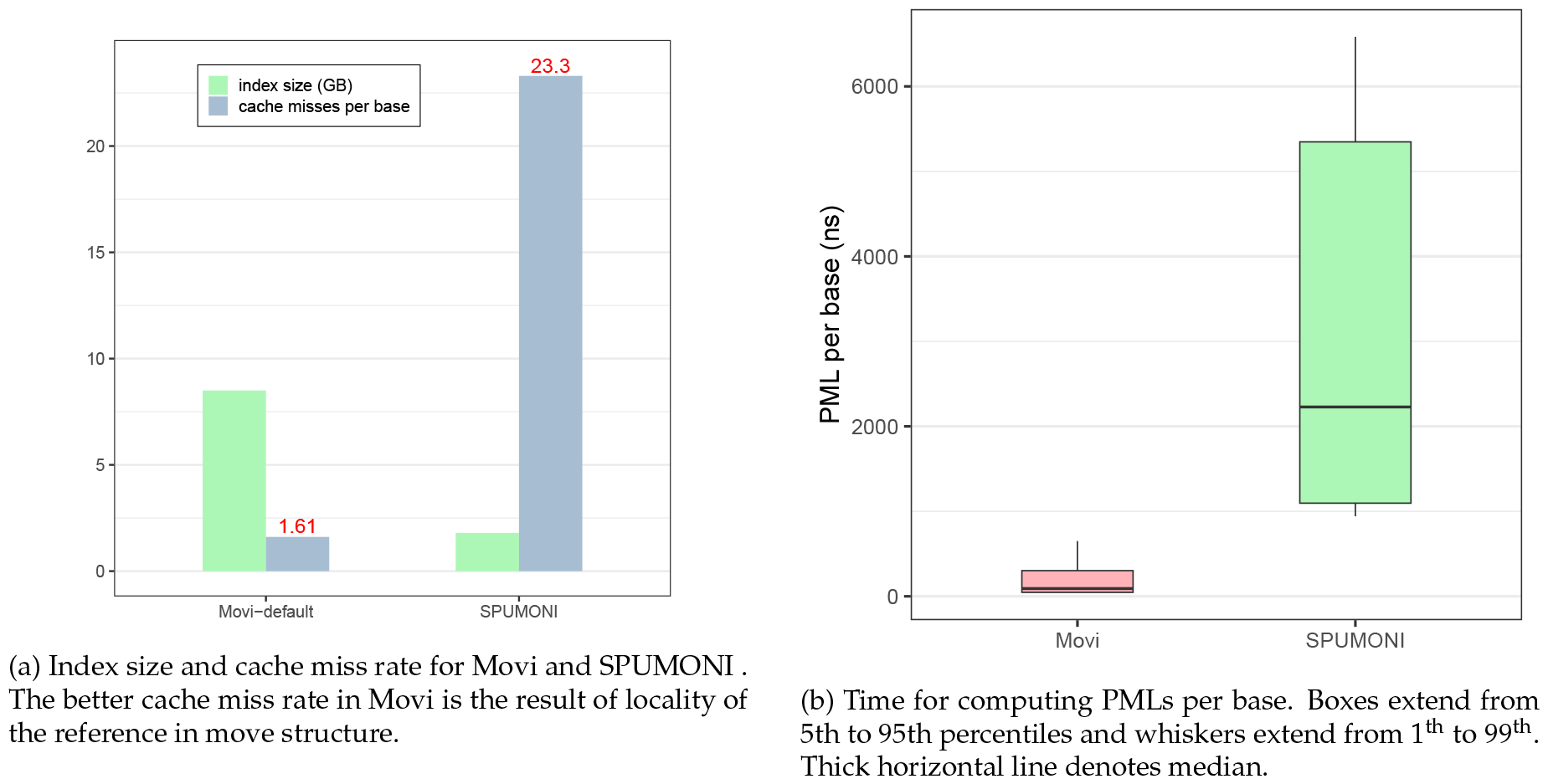
Comparisons of Movi-default and SPUMONI in terms of query speed and predictability.

To assess the latencies of `LF`-mapping executed by SPUMONI and Movi more precisely, we employed the chrono high-resolution clock in C++ to make nanosecond-level latency measurements for their inner loops. The distribution of these latencies is visualized as boxplots in Figure 3b. We observed that iterations of the Movi inner loop were about 11.5 times faster than those of SPUMONI (comparing means). The 99th-percentile latency observed for Movi’s inner loop (650 ns) was smaller than the 1st-percentile latency observed for SPUMONI’s inner loop (942 ns). The median latency observed for Movi’s inner loop (91 ns) was also much smaller than SPUMONI’s (2,228 ns). Note that a single last-level cache miss is roughly thought to take 100 ns, or 300 clock cycles on a 3 GHz processor.

Besides variability in inner loop performance due to cache misses, we measured the number of fast-forward iterations and repositioning scans in each of Movi’s modes. These were discussed in Section 2.3. As expected, the number of operations was bounded by a small constant for Movi-constant. For Movi-default, the number of operations varied much more, as seen in Figure 4. Detailed statistics are presented in supplementary materials Table S3. While we earlier observed that Movi-default was faster than Movi-constant on average, here we saw that Movi-constant’s inner loop performed a smaller and more predictable number of operations, which is advantageous in situations where the algorithm must keep up with the output of an instrument in real-time. However, the average number of fast-forwards performed in Movi-default’s loop compared to Movi-constant’s was only about about 1.2 times greater, and the average number of repositioning scans was only about 2.5 times greater. The fact that Movi-default is still faster than Movi-constant despite this difference is likely because of the fact that Movi-constant requires a larger index, which in turns incurs more cache misses overall.

**Figure 4:**
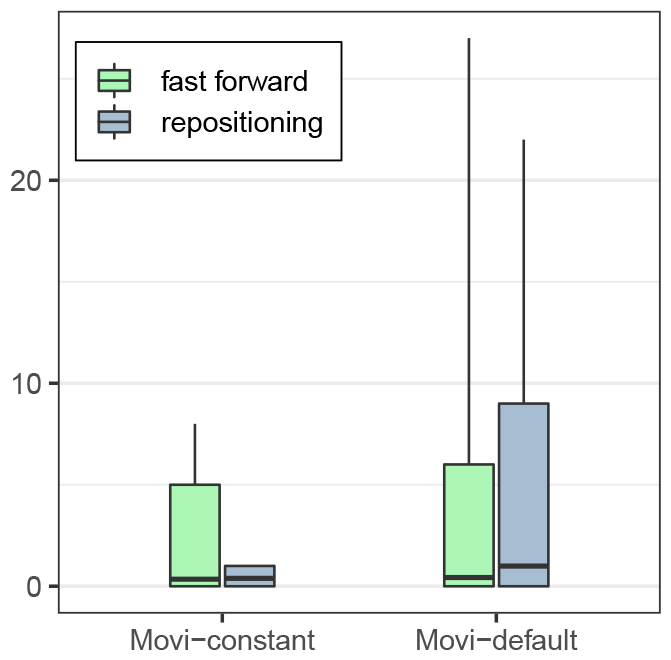
The number of fast-forwards and repositioning scans in each mode of Movi. Movi-constant is guaranteed to use a constant number of memory accesses per `LF`-mapping. Boxes extend from 1st to 99th percentiles and whiskers extend from 0.1^th^ to 99.9^th^. Horizontal line denotes mean. In all cases, the median is 0.

### 3.3 Extrapolation to nanopore throughputs

Using per-base speeds measured for the Zymo input data (presented in Table 1), we extrapolate to measure their ability to analyze nanopore sequencing data in a real-time adaptive sampling context. We assume that the sequences are base-called immediately. Considering that the sequencing speed of each nanopore of an Oxford Nanopore (ONT) instrument is 420 base pairs per second, SPUMONI ‘s speed is sufficient to simultaneosuly handle 904 channels (pores) at once. On the other hand, Movi can handle 26,890 simultaneous channels, surpassing the total number of channels in the largest flow cell available for the PromethION device: 2,675 channels^4^. Assuming perfect linear scaling, about 5 simultaneous Movi threads (each handling 16 reads concurrently) would be sufficient to handle the aggregate output of 48 PromethION flowcells.

### 3.4 Backward search for count queries

Besides pseudo-matching lengths, another full-text query is the “count” query, which reports the number of distinct locations where the query occurs as a substring of T. A count query involves a sequence of backward-search steps, each step using one additional character of the query.

In Movi, backward search begins by finding the range of BWM rows that have the final (rightmost) query character as a prefix. In subsequent steps, `LF`-mapping-like steps are used to advance this range’s top and bottom pointers to additionally match the next query character to the left (i.e. a longer suffix of the query), obtaining an the interval of BWM rows beginning with the longer suffix. This repeats until the query is exhausted or until the range becomes empty, indicating that the query does not occur. In Movi, updating the top and bottom pointers is exactly analogous to the repositioning procedure described in Methods 2.3, except that the choice of j^up^ or j^dn^is determined by whether we are updating the top pointer (in which case we use j^dn^) or the bottom pointer (in which case we use j^up^).

To measure backward search performance, we used Mason [18] to simulate 10 million 150-bp unpaired reads from a FASTA file containing the complete genomes of the 7 bacterial species in the Zymo community, which was also used for Results section 3.1. We generated error-free reads to ensure that backward search would iterate over all query characters. We compared Movi’s efficiency to that of r-index, which supports the same query. Note that SPUMONI does not support this same query. We observed that r-index took 44m:05s, while Movi took 2m:43s, a 16-fold improvement. On the other hand, the Movi-default index was about 3 times larger than the r-index, consistent with other results showing the move structure to be larger.

**Table 2:**
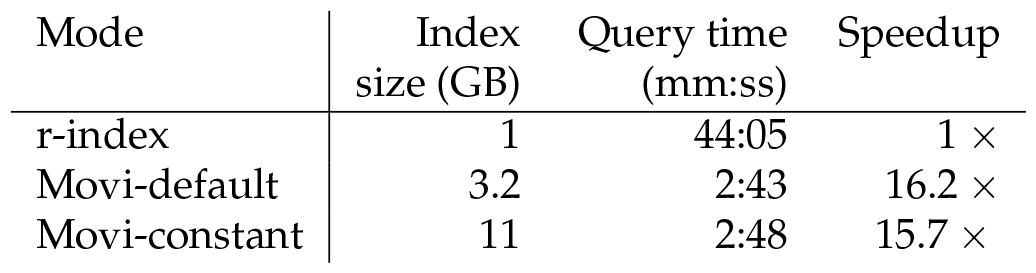
Time and index size required to execute the count query with Movi and the r-index. Both default and constant modes of Movi are very fast while the constant mode uses more memory, because it stores the repositioning pointers in each row.

### 3.5 Scaling to human pangenomes

We next evaluated the scalability of Movi using human genome haplotype assemblies from the Human Pangenome Reference Consortium (HPRC) [15]. We selected various numbers of haplotypes, ranging from 1 to 94, which includes all available haplotypes. We measured the overall size and scalability of Movi’s indexes (based on the move structure) when compared to SPUMONI (based on r-index) and Fulgor (based on colored compacted de-bruijn graph). Note that Fulgor’s index also stores “color” information (associating k-mers with haplotypes), which is not a type of information stored in the Movi or SPUMONI indexes. We used k=31 and m=19 when building the Fulgor indexes.

We measured each tools’ ability to scale to larger pangenomes in Section 3.5. As a baseline for measuring scalability, we reported the number of distinct k-mers in the input according to Fulgor’s `stats` command (“kmer-count” column). As a second baseline, we also reported the number of runs in the BWT according to Movi (“r” column). As seen in Figure 5, the size of the 94-haplotype indexes were less than 2 times the size of the 5-haplotype indexes for all three tools. Movi exhibited the best scaling factor, with its 94-haplotype index using about 1.2 times the space as its 5-haplotype index. The 94-haplotype index for Fulgor and SPUMONI used 1.38 and 1.86 times the space as their 5-haplotype indexes respectively. This highlights the advantages of compressed indexes, including full-text indexes, when indexing large pangenomes.

**Figure 5:**
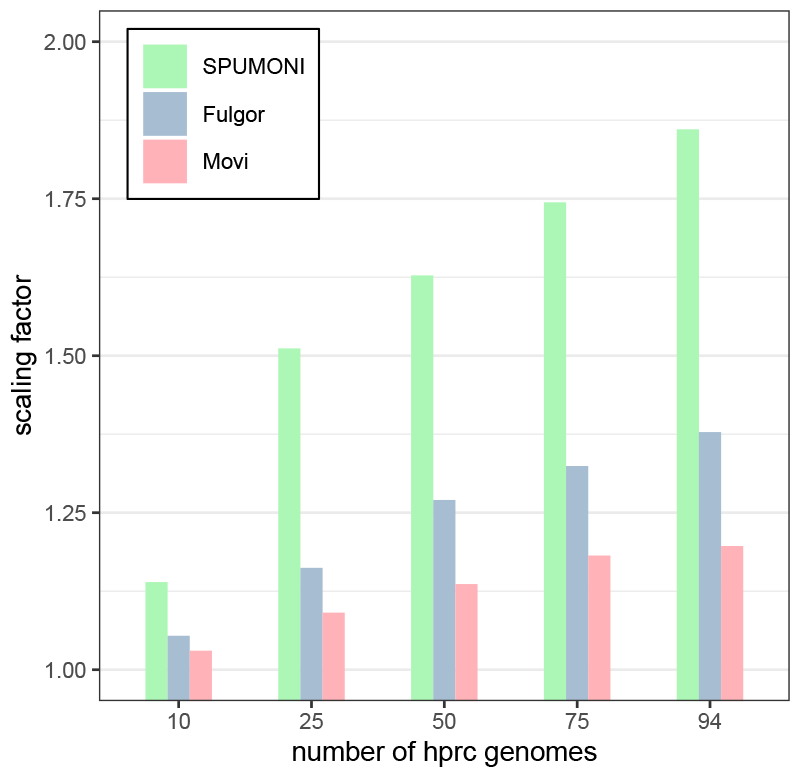
The scaling factor is computed by dividing the size of each tool’s index by the size of the index of that tool built over 5 HPRC genomes. All tools have small scaling factors for pangenomes. While Movi’s index is the largest compared to the other two, it has the best scaling factor for any number of hrpc genomes.

We also observed that the size of Fulgor’s index was considerably smaller than both SPUMONI ‘s and Movi’s. Fulgor’s index includes both k-mer mapping and color class information, i.e. information about which k-mers occur in which haplotypes. In this experiment, there are relatively few colors and so color-class information makes up a smaller portion of the index. Running Fulgor’s ‘stats’ command on indexes created in Section 3.5 showed that between 1% to 5% of the index is dedicated to color information.

We also evaluated query speed for each tool using a simulated long read sample and a “combined” sample, consisting of both simulated reads and real reads from a human gut sample. This allows us to measure performance in a scenario where many input reads do not have a long match to the reference pangenome. The results are shown in Table S5 and are similar to those presented in Section 3.1, with Movi being fastest followed by Fulgor and SPUMONI.

**Table 3:** Indexes are built over different number of HPRC assemblies: 1, 5, 10, 25, 50, 75, 94 (all)

| reference | fasta (GB) | kmer-count ($\times 10^9$) | Fulgor (GB) | r ($\times 10^9$) | SPUMONI (GB) | Movi (GB) |
| --- | --- | --- | --- | --- | --- | --- |
| HPRC 1 | 2.9 | 2.50 | 3.1 | 3.33 | 6 | 62 |
| HPRC 5 | 15 | 2.70 | 3.7 | 3.53 | 8.6 | 66 |
| HPRC 10 | 29 | 2.79 | 3.9 | 3.65 | 9.8 | 68 |
| HPRC 25 | 74 | 2.94 | 4.3 | 3.84 | 13 | 72 |
| HPRC 50 | 174 | 3.06 | 4.7 | 4.02 | 14 | 75 |
| HPRC 75 | 214 | 3.13 | 4.9 | 4.14 | 15 | 78 |
| HPRC 94 | 268 | 3.19 | 5.1 | 4.24 | 16 | 79 |

## 4 Discussion

We introduced Movi, a cache-efficient, scalable tool for pangenomic indexing and read classification. Movi’s index is based on the move structure which is a full-text index with a scaling factor superior to competing approaches like SPUMONI and Fulgor. Movi is extremely fast, due both to its excellent locality of reference which in turn minimizes cache misses, and to our novel strategy for hiding the remaining cache-miss latency by processing many reads concurrently. Movi’s rapid and predictable query speed makes it well suited to applications like nanopore adaptive sampling. Movi can process the base-called output of a fully loaded PromethION using 12 threads.

The move structure’s simple tabular structure suggests simple ways to partition and distribute it across nodes of a computer cluster while minimizing inter-node communication. It can simply be divided into separate, contiguous chunks of rows, which can then be distributed. Execution of a pattern-matching query will require some jumps between nodes (i.e. a longer-distance LF query), but will frequently require only sequential or nearby jumps (fast-forwards and repositions) that do not require moving across nodes. This provides a much more favorable substrate for distributed computing compared to r-index, which is characterized by complex and unpredictable memory accesses.

Another key advantage of our full-text indexing approach is that it does not require the user to select any key parameters ahead of time. This is in contrast to k-mer based or minimizer-based approaches, for which the user must be aware of the potential pitfalls of choosing suboptimal parameters.

A limitation of Movi is the fact that the M table is large compared to all the other tools assessed here (besides minimap2). In the future, it will be important to reduce the footprint of Movi’s index. This could be accomplished, for instance, by adopting the minimizer digestion strategy of SPUMONI 2 [6]. Another space-saving measure could be to losslessly compress the move structure using, e.g., the columnar compression strategies investigated by Brown et al. in 2022 [11].

We also hope to expand Movi’s applicability to more query types. For instance, Movi could be adapted to handle multi-class classification by augmenting the index with suffix array or “document” information[19].

While Fulgor [20] optimizes space and time by capitalizing on long unitigs and explicitly storing the corresponding strings, we can adopt a similar strategy by leveraging substructures within the BWT. One such approach is to enhance query efficiency by reordering the BWT rows. This technique can be seamlessly integrated into Movi, enabling further cache efficiency and greater speed. By incorporating reordering, Movi has the potential to achieve even greater query performance.

## Acknowledgements

This work was supported by NIH grant R01HG011392 to BL and NSF-135491. NKB and TG were supported by NSERC grant RGPIN-07185-2020 to TG. NKB was also supported by a Johns Hopkins University Computer Science PhD Fellowship.

## Author contributions

MZ and BL designed the method, with help from NKB, OYA, and TG. MZ wrote the software with help from NKB. MZ performed the experiments. All authors contributed to the manuscript.

## Supplementary Materials

### Algorithm S1: The fast forward algorithm, “ff”

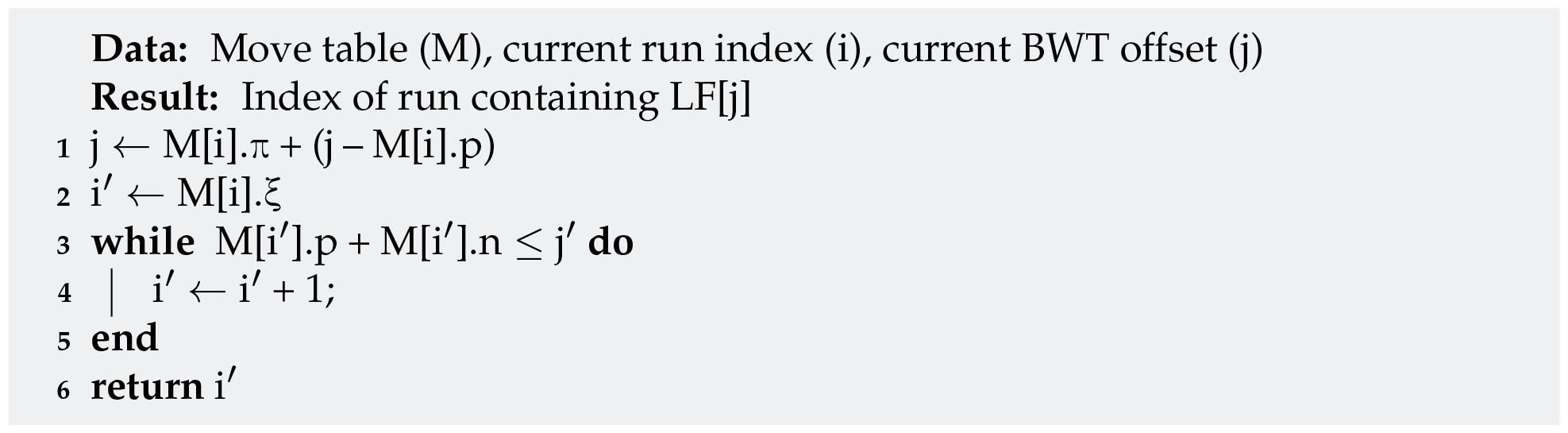

### Algorithm S2: PML computation using move structure. RepositionUp and RepositionDown are performed using scanning in the default mode, or the explicit pointers in the constant mode.

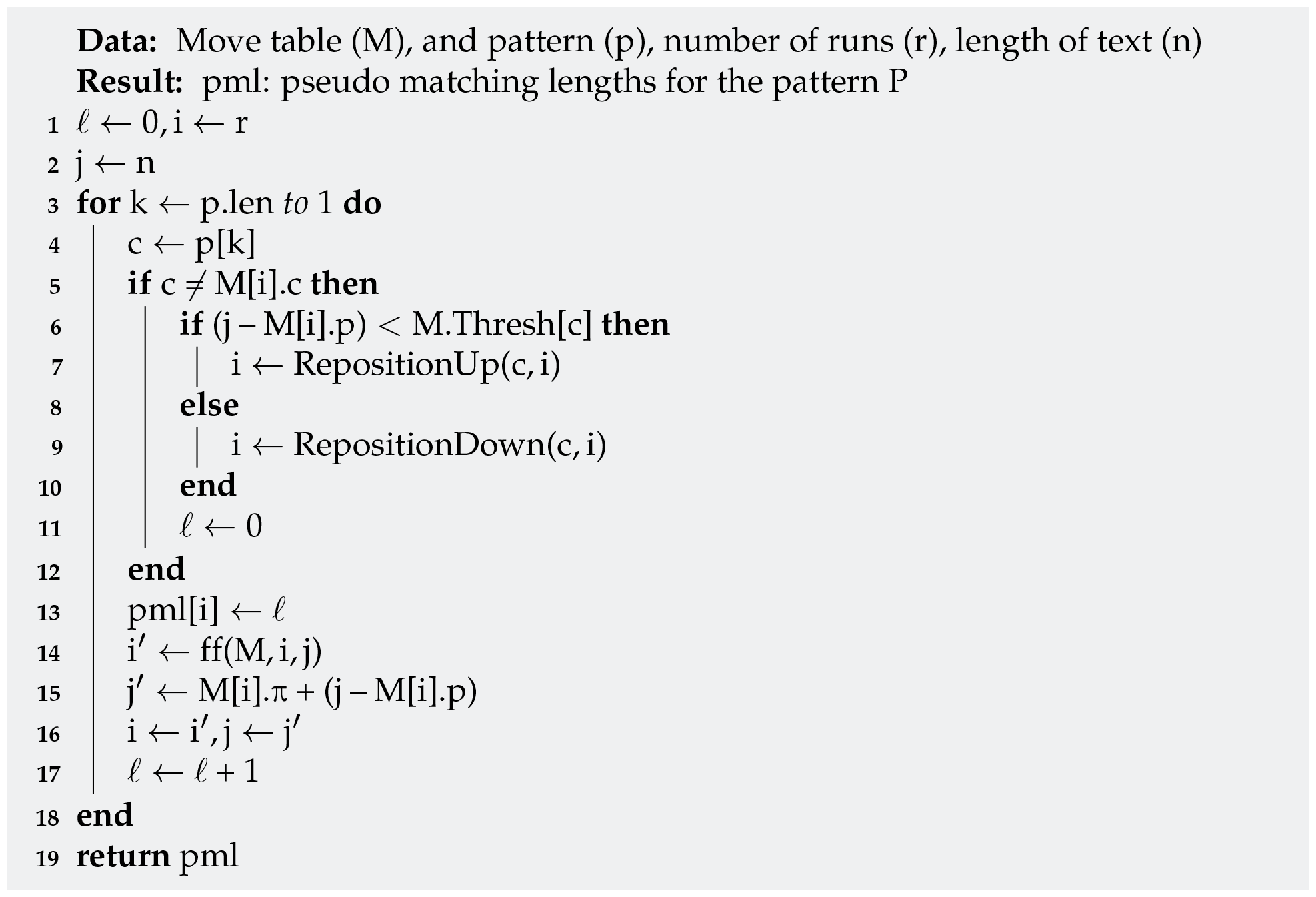

### Algorithm S3: The prefetching algorithm in move structure for computing PMLs. Each read is assigned to a class called “Strand” to be processed.

After all the PMLs for a read in one Strand is computed, the Strand is updated to process the next read, “NextRead” retrieves the next read from the input file and assigns it to a strand. “NextPML” generates the pseudo matching length for the next base in the read assigned to strand s. “Prefetch” triggers an asynchronous retrieaval of the memory containing the destination row of the `LF`-mapping, which will be used later.

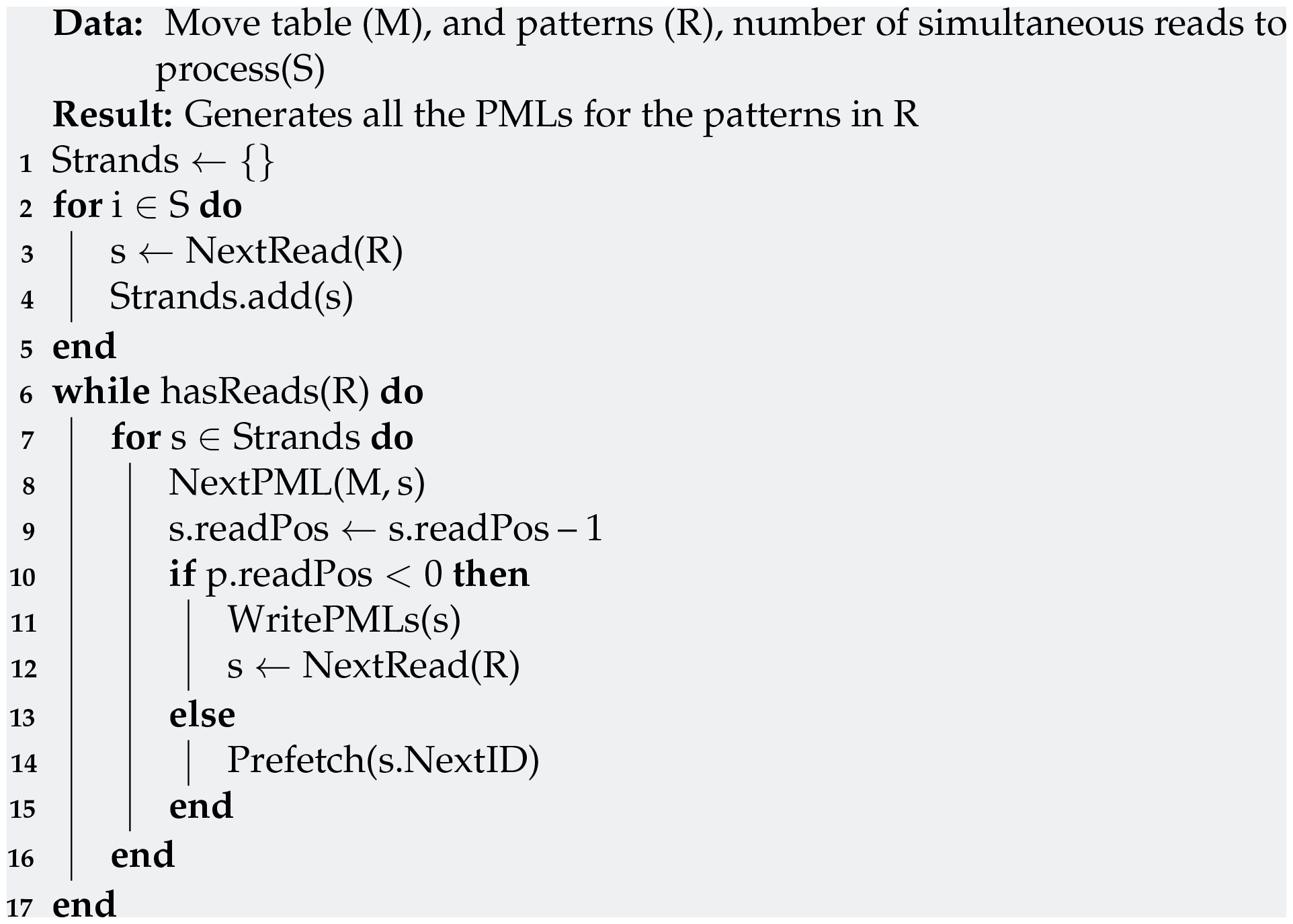

**Figure S1:**
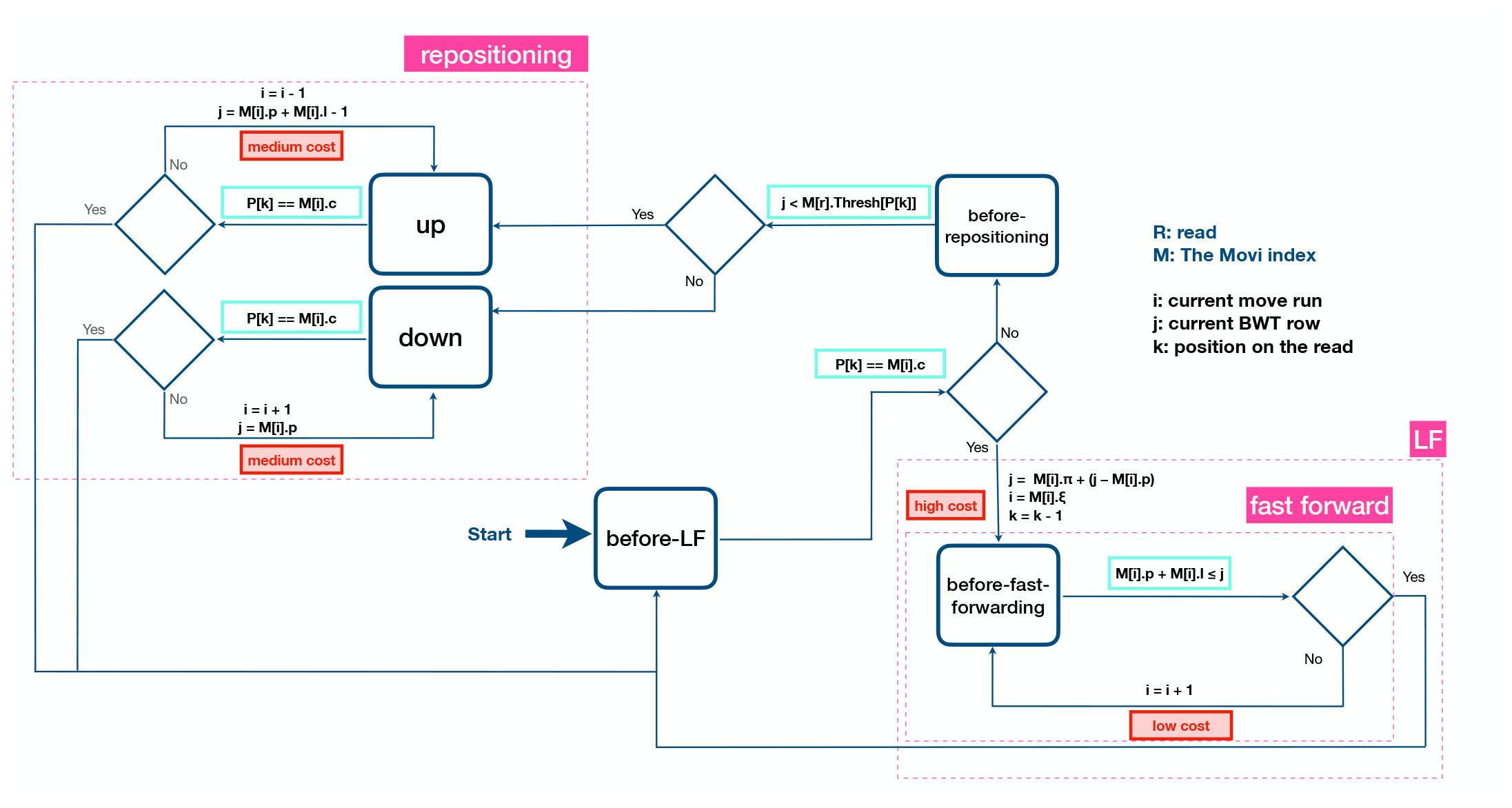
Full schematic of the pseudo matching length (PML) computation with Movi-default. For Movi-constant pointers and direct-address lookups are used instead of the linear-scan loops shown in the figure.

**Table S1:**
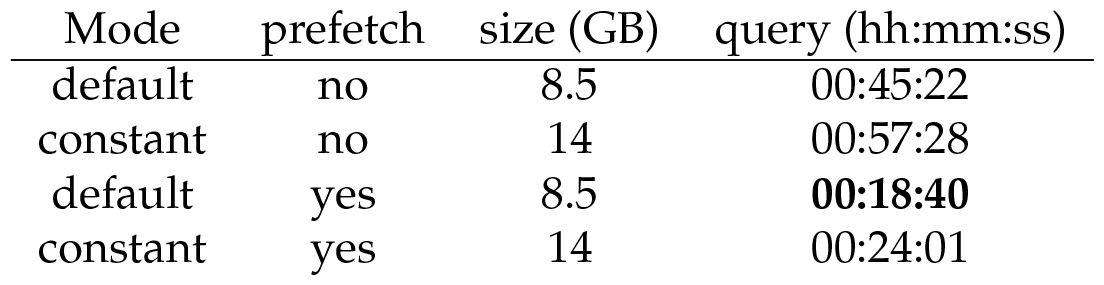
Comparison of different Movi modes. “Constant” refers to the mode that uses both (a) Nishimoto and Tabei’s splitting procedure for ensuring a constant number of fast forwards and (b) constant-time direct-address lookups in the repositioning procedure. “Default” mode does not use splitting and uses linear scans in the repositioning procedure. “Prefetch” refers to whether Movi is using its latency-hiding approach. When “prefetch” is “yes,” Movi is processing 16 reads concurrently and using prefetch instructions to hide cache-miss latency.

**Figure S2:**
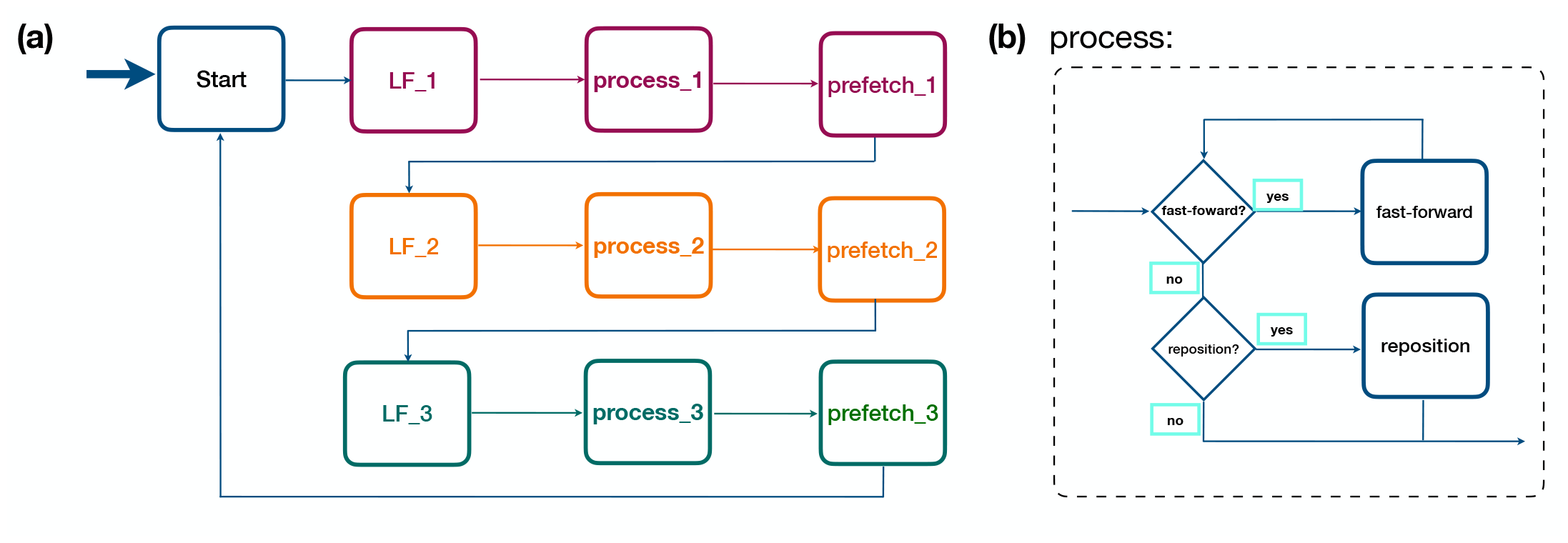
Computing PMLs with prefetching in Movi. (a) Shows how each read is processed until the LF step which is the highest cost. Then a prefetching step is triggered to fetch the memory required to do the LF. While the memory is being prefetched, the processor moves to processing another read for which the memory required is already prefetched, (b)The process which is performed after the LF for each read, note that all the steps in this processa are low cost or medium cost, therefore, these are much faster compared to the LF step for which the memory is prefetched.

**Figure S3:**
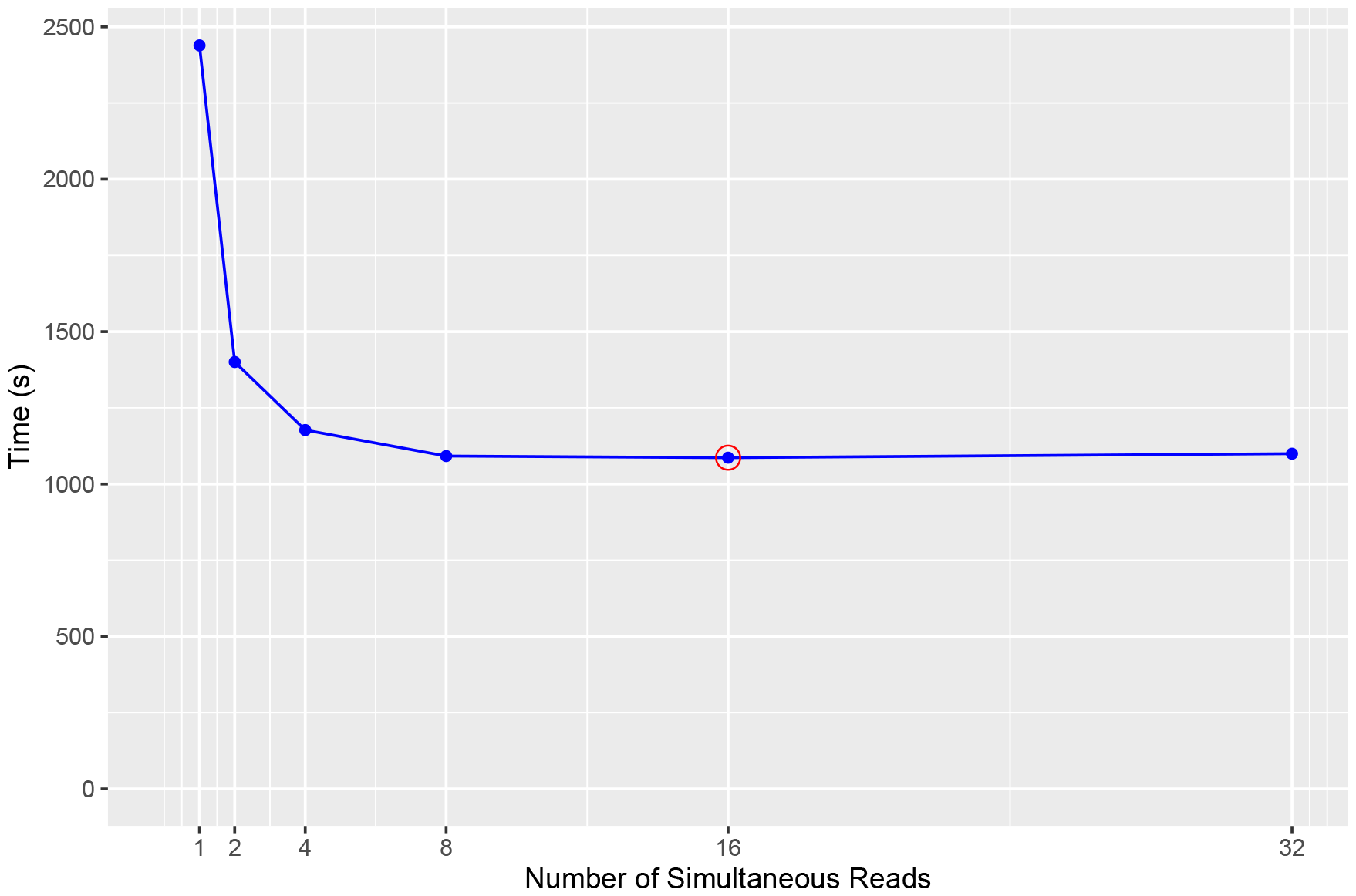
Time required to process the reads from the Zymo community using the latency-hiding strategy as a function of the number of reads being processed concurrently. The latency-hiding benefit accrues rapidly up to 8 threads, then plateaus. Movi processes 16 concurrent reads by default (red circle).

**Figure S4:**
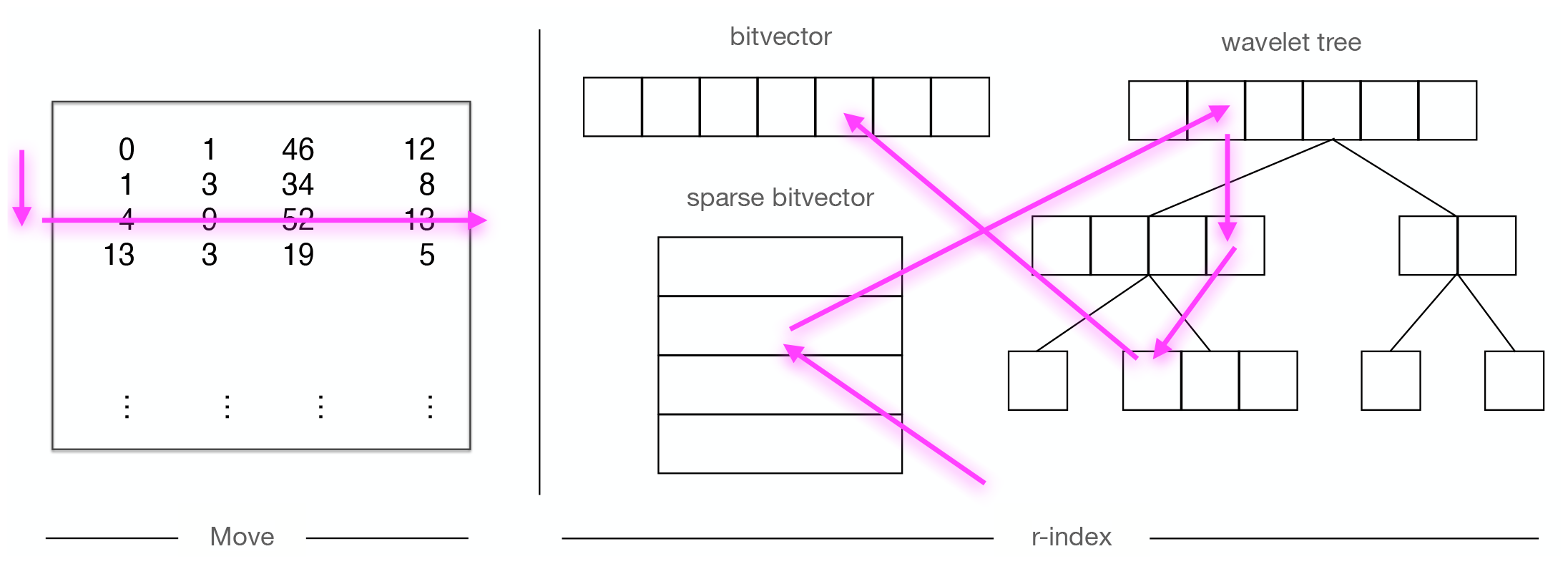
Approximate schematic for illustrating how memory accesses are induced for `LF`-mapping in r-index and move structure. This is not an exact representation of how queries work in these indexes and is just intended to roughly show the memory access patterns.

table for comparing the cache misses between movi and spumoni similar statistics are shown in sub-fig:cache

**Table S2:** cache miss comparison.

| Tool | index (GB) |  | speed (10<sup>6</sup> base per second) |  | cache miss per base |  |
| --- | --- | --- | --- | --- | --- | --- |
| Movi | 8.5 | x4.7 | 11.29 | x 30 | 1.61 | - |
| SPUMONI | 1.8 | - | 0.38 | - | 23.30 | x 14.47 |

**Table S3:** Fast-forward and repositioning statistics for the Zymo sample.

| operation | mode | mean | sd | max |
| --- | --- | --- | --- | --- |
| fast-forward | Movi-default | 0.4189303 | 3.682711 | 7694 |
|  | Movi-constant | 0.3395261 | 0.913855 | 9 |
| repositioning | Movi-default | 1.039042 | 2.416862 | 4345 |
|  | Movi-constant | 0.4068258 | 0.4912419 | 1 |

**Table S4:**
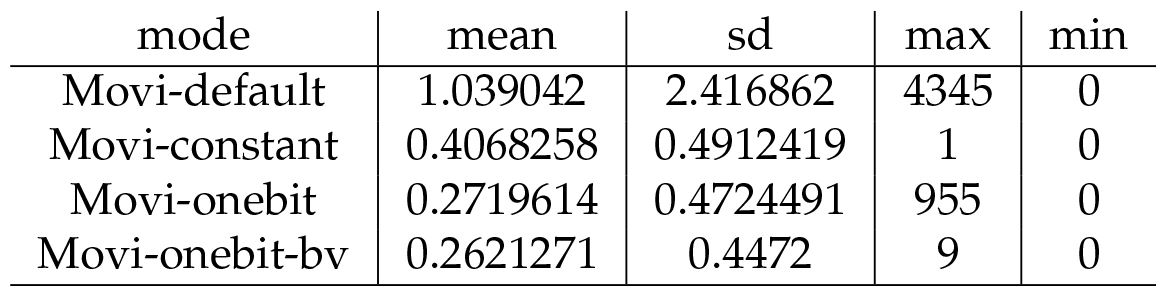
Scans statistics.

**Table S5:** Query speed for the hprc dataset. The simulated sample consists of long reads simulated by PBSIM2[21] from a human genome. The combined sample consist of both the simulated reads and a human gut metagenomic sample (`SRR9847854`). The results for Movi in this table are evaluated without the latency hiding strategy.

| sample | reference | Fulgor (hh:mm:ss) | SPUMONI (hh:mm:ss) | Movi (hh:mm:ss) |
| --- | --- | --- | --- | --- |
| simulated (350K) | hprc 1 | 00:19:30 | 02:28:13 | 00:09:58 |
|  | hprc 94 | 00:22:39 | 02:38:12 | 00:17:19 |
| combined (11M) | hprc 1 | 01:50:19 | 26:51:22 | 01:35:37 |
|  | hprc 94 | 02:04:21 | 28:31:58 | 01:44:17 |

### Commands used for running the tools

~~~
Building The indexes:
 $ spumoni build -i <reference_files_list> -o <index_prefix> -P -n
 $ fulgor build -l <reference_files_list> -o <output_prefix> -k 31 -m 19 -t 16
 $ minimap2 -x map-ont -d <index_file> <reference_file>
 $ bowtie2 --large-index <reference_file> <index_prefix>
Running the queries:
 $ spumoni run -r <index_prefix> -p <reads_file> -P -n
 $ fulgor -i <index_file> -q <reads_file> -o <output_file> -t 1
 $ minimap2 --secondary=no -t 16 -x map-ont <index_file> <reads_file> -o <output_file>
~~~

## Footnotes

1 https://github.com/maxrossi91/pfp-thresholds

2 https://github.com/drnatebrown/r-permute

3 https://valgrind.org/docs/manual/cg-manual.html

4 https://nanoporetech.com/products/specifications

